# Colorful connections: pigment-based plumage and breeding condition are associated with gut microbiome variation in the Common Yellowthroat

**DOI:** 10.64898/2026.06.16.732689

**Authors:** Alix E. Matthews, Maxima Gomez-Palmer, Sebastian Gallego, Mads Moore, Lan-Nhi Phung, Daniel T. Baldassarre, Marcella D. Baiz

## Abstract

Carotenoid- and melanin-based plumage coloration traits are key signals in avian communication and sexual selection as they are often thought to provide “honest” information about individual condition and fitness. These traits arise through distinct but interconnected physiological and genetic pathways. Recent work suggests that there may be a link between host-associated gut microbiota and the functional pathways leading to pigment-based plumage coloration, but this remains largely unexplored in wild populations. To address this gap, we tested whether variation in plumage coloration, as well as breeding condition, is associated with gut microbiome variation in wild populations of male Common Yellowthroats (Parulidae: *Geothlypis trichas*). We quantified multiple plumage coloration traits and characterized gut microbiome bacterial diversity using 16S rRNA metabarcoding. Through a comprehensive modeling framework, we found that individuals with brighter, more orange-tinted breast feathers and smaller cloacal protuberances (a proxy for breeding condition) exhibited higher gut microbiome diversity. At the taxonomic level, *Methylobacterium-Methylorubrum*, a carotenoid-producing bacteria, showed strong associations with multiple plumage traits, including mask area, breast feather hue, and saturation. Our results demonstrate that gut microbiome diversity is associated with variation in carotenoid-based coloration traits and breeding condition in Common Yellowthroats. More broadly, these results highlight the potential for host-microbiome interactions to shape phenotypic variation through physiological pathways in wild animal populations.

## INTRODUCTION

Sexual selection has played a central role in the evolution of conspicuous traits used in mate choice and intrasexual competition across animals (Andersson 1994). Theory predicts one way that sexual selection can favor secondary sexual traits (e.g., ornaments, armaments, coloration, behavioral displays) is when they signal reliable information about individual quality. Consequently, many secondary sexual traits are thought to function as “honest signals” of individual condition. Whether honest signals arise through condition-dependent production costs (Zahavi 1975, Kodric-Brown and Brown 1984) or via intrinsic links to physiology (Hill 2011, Hill et al. 2023) remains a subject of debate. Nevertheless, carotenoid (e.g., red, orange, yellow) and melanin (e.g., black, brown)-based plumage traits in birds are among the best studied examples of such signals and are linked to key components of individual fitness including reproductive success (Bulluck et al. 2017, Slevin et al. 2019) and survival (Hõrak et al. 2001, Rolland et al. 2024). Understanding the mechanisms that contribute to these signal-based phenotypes is therefore critical for understanding how selection acts on them in natural populations.

Carotenoid- and melanin-based coloration arises through distinct but interconnected physiological pathways. On one hand, birds (and most other animals) cannot synthesize carotenoids *de novo*, so they must acquire these pigments from external sources and then process them internally (Toews et al. 2017). On the other hand, melanin pigments are synthesized endogenously (D’Alba and Shawkey 2019). Once acquired or synthesized, the deposition of these pigments into external integuments (e.g., feathers) to produce coloration and patterning is shaped by multiple processes including pigment transport, metabolic reaction efficiency, and oxidative stress (Galván and Alonso-Alvarez 2009, LaFountain et al. 2015, McNamara et al. 2021). These physiological processes can be impacted by hormones; for example, as birds enter breeding condition, sex hormone (e.g., testosterone) levels fluctuate (Moore et al. 2005) and can influence both oxidative stress and immune function, potentially altering pigment processing and deposition (Folstad and Karter 1992, Casagrande et al. 2012). Additionally, hormones can interact with genetics to impact pigment-based ornamentation by regulating the expression of genes that control coloration (Khalil et al. 2023). Genes that control avian plumage coloration have been identified by examining differences between species, and some genes have strong effects on plumage traits. For example, agouti signaling protein (*ASIP*) controls the presence or absence of melanin traits in predictable (i.e., Mendelian) ways (Toews et al. 2016, Baiz et al. 2020). β-carotene oxygenase 2 (*BCO2*) and others (e.g., *CYP2J19*, *BDH1L*) are implicated in carotenoid processing (Walsh et al. 2012, Lopes et al. 2016, Toews et al. 2017, Baiz et al. 2021, Bennett et al. 2025). However, many pigment-based traits tend to show continuous variation within a species (Baldassarre et al. 2022, Sly et al. 2022). Thus, the mechanisms identified to date likely represent only a subset of the physiological and genetic processes that generate the substantial variation in pigment-based plumage traits within populations. One potentially important, yet underexplored, contributor to this variation is the gut microbiome, which can influence nutrient availability, metabolism, and host physiology in ways that may mediate pigment expression.

The gut microbiome has been described as a “second genome” that shapes host nutrition, immune function, and metabolic processes (Zhu et al. 2010). Microbes can alter the availability and transformation of dietary compounds (Roggenbuck et al. 2014, Maccaro et al. 2024), shape energetic and metabolic pathways (Hammer et al. 2023, Houtz et al. 2023), and modulate host immunocompetence and stress (Houtz et al. 2022, Slevin et al. 2026), all of which are processes that are linked to individual fitness. These functions suggest multiple potential routes by which the gut microbiome could influence pigment-based ornament expression in hosts. Consistent with this possibility, a growing body of literature has revealed relationships between gut microbiome variation and coloration traits in both vertebrate (Nguyen et al. 2020) and invertebrate (Koellsch et al. 2024) systems. A significant amount of work to date in this area has been conducted in avian systems, likely because plumage coloration traits are highly variable, measurable, and tightly linked to sexual selection. For example, in a natural experiment involving two hybridizing warblers, Baiz et al. (2024) found that admixed individuals with yellower plumage (based on a scoring system of nine plumage characters) had higher relative abundances of several known carotenoid-producing bacterial taxa in their gut microbiomes, as well as higher abundances of *Kyoto encyclopedia of genes and genomes* (KEGG) orthologues and enzymes involved in carotenoid biosynthesis pathways based on predictive metagenomics. Similarly, other studies have reported associations between gut microbiome variation and both carotenoid- and melanin-based traits, such as beak brightness and mask darkness (Marques Silva et al. 2023, Slevin et al. 2025). Complementing these correlational studies, disruption of the gut microbiome in an experimental setting (i.e., through antibiotic treatments) has been shown to alter the deposition of carotenoids into feathers (Lind et al. 2021). Together, these studies demonstrate a growing connection between the host (specifically avian) gut microbiome and secondary sexual traits. Furthermore, gut microbiome diversity has been positively associated with testosterone levels (Escallón et al. 2017, Houtz et al. 2025), suggesting that the physiological shifts that accompany breeding and sexual selection are also linked to the microbiome. However, it remains unclear how consistent these patterns are within and across wild populations and which, if any, microbial taxa specifically drive these relationships.

Wood-warblers (Parulidae) are an excellent model for investigating these connections because they are a highly ecologically and phenotypically diverse clade. They are among the most colorful bird families in the Americas and exhibit substantial inter- and intraspecific variation in their pigment-based plumage coloration and patterning (of which many traits are quite prominent; Stephenson and Whittle 2013). As such, their phenotypic traits have been extensively studied in the context of sexual selection, particularly during the breeding season (Thusius et al. 2001b, Dunn et al. 2008, Freeman-Gallant et al. 2010, Jones et al. 2017). Links between parulid gut microbiome variation and plumage have been made in the hybridizing *Vermivora* genus; however, plumage coloration was characterized by a subjective scoring scheme and was confounded by genetic ancestry (Baiz et al. 2024). This limits our ability to determine how variation in the gut microbiome relates to fine-scale differences in plumage traits under sexual selection within populations, or to disentangle those relationships from the physiological changes associated with breeding condition.

In this study, we examined the relationships between gut microbiome variation, pigment-based plumage traits, and breeding condition within a single, widely distributed migratory North American parulid, the Common Yellowthroat (*Geothlypis trichas*). Male Common Yellowthroats exhibit two conspicuous pigment-based traits: a carotenoid-based yellow “bib” (throat, breast, and belly) and a melanin-based black facial mask, both of which function as “honest” signals of individual quality and are implicated in sexual selection (Dunn et al. 2008, Sly et al. 2022). Our overarching goals were to (1) measure the extent to which plumage traits and breeding condition are associated with variation in the gut microbiome and (2) identify specific microbial taxa associated with those traits in this species. To address these goals, we quantified carotenoid- and melanin-based plumage traits and breeding condition (indexed by cloacal protuberance height) in male Common Yellowthroats and characterized their gut microbiome diversity and composition. By linking multiple quantitative plumage phenotypes and an index of breeding condition with both community- and taxon-level microbiome data, our approach allowed us to model how microbiome variation covaries with sexual traits in natural populations.

## METHODS

### Study Species, Field Sites, and Sample Collection

We captured Common Yellowthroats in 2022 and 2023 between May and June during the breeding season in eastern North America (Pennsylvania [2022, 2023] and New York [2023], USA), where they are short- to long-distance migrants. Common Yellowthroats are a widely distributed species that occurs in dense, low-growing vegetation most often in wet areas (Stewart 1953, Guzy and Ritchison 2020). They forage low to the ground for invertebrate prey, of which Diptera, Coleoptera, and Lepidoptera are prominent (Miller et al. 2025).

To quantify gut microbiome composition and diversity, we collected fecal samples to conduct 16S rRNA metabarcoding. To do so, we lured individuals for capture in a mist net using song playback (n = 56 males). Upon capture, individuals were placed in a clean paper bag for ten minutes. After this, we removed the bird and collected feces from the bag by scraping it directly into a sterile tube containing 500µL of 0.5% SDS buffer (White and Densmore 1992). Fecal samples were initially stored at ambient temperature in the field for up to two weeks, after which we transferred them to a −20°C freezer until DNA extraction.

Morphometric data were then collected and breeding condition was assessed before affixing a uniquely numbered United States Geological Survey aluminum leg band for individual identification. To assess breeding condition, we scored the size of the cloacal protuberance based on height from 1 to 3, where 1 (low) indicates non-breeding condition and 3 (high) indicates full breeding condition (Pyle 2022).

After a fecal sample was collected and other data recorded, we plucked 5-10 breast feathers for reflectance spectrometry and captured a series of photographs for quantitative size measurements of mask and bib plumage ornaments. We held individuals against a standard paper grid of 1cm x 1cm black-and-white checkered squares and captured two photographs per left and right side of the head and two photographs of the ventral side (Figure S1).

### Reflectance Spectrometry

To measure plumage reflectance, we mounted five breast feathers on black matte cardstock in an overlapping pattern to create a feather patch similar to their natural position on the bird’s breast. We used an Ocean Optics Spectrometer (SR4-UV400-25) with a Pulsed Xenon Light Source (PX-2) and a reflection probe (QR400-7-SR-BX) to measure reflectance in the ultraviolet (UV; 220-400nm) and visible (400-750nm) spectra. Using a reflection probe holder that excluded all ambient light, we held the probe over the feathers at a 45° angle to take measurements. The probe extended 27mm into the block (i.e., distance from the probe tip to the block’s exit opening, per manufacturer recommendations). We used OceanView version 2.0.20 to record reflectance data. Measurements were calibrated between each sample with a diffuse white reflectance standard (WS-1) and a dark standard (black matte cardstock) to account for background noise. For each sample, we measured reflectance three times by positioning the probe over three different regions of the mounted feathers. We used the R package “pavo” version 2.9.0 (Maia et al. 2019) to average the three spectra and to smooth (span = 0.1; to reduce noise) the reflectance curves before further colorimetric analyses.

Following spectral pre-processing, we estimated photoreceptor quantum catch values for each of the four avian single cones and the double cone using the Eurasian Blue Tit (*Cyanistes caeruleus*) visual model and *vismodel* function. Values were calculated under a D65 standard daylight illuminant and scaled as relative quantum catches. We then projected the quantum catch values from the single cones into tetrahedral color space using the *colspace* function to obtain chromatic coordinates for each sample. We extracted hue (h.theta) and saturation (r.achieved) from the color space, and used the double cone stimulation as our measure of brightness (lum). For hue, theta (h.theta) and phi (h.phi) were correlated (R = -0.77, p < 0.001), so we only used theta for downstream analyses. We used these variables to quantify variation in feather coloration as perceived by the avian visual system.

### Ornament Size Measurements

Ornament (i.e., black mask and yellow bib) size was estimated from photographs of captured birds. All images were imported into ImageJ (https://imagej.net/ij/) and scaled before each measurement to the 1cm x 1cm background grid. To measure mask size, we traced the outline of the mask. To measure the bib area, we used the “threshold” plugin to delimit the extent of the bib with the following settings: hue to 20-50, saturation to 100-255, and brightness was increased from zero until the image reached the edge of the bib. Extraneous pixels were removed. Ornament size estimates were averaged across replicate images from each ornament for downstream analyses. All size measurements were made by the same individual for consistency and double-checked by two others.

### DNA Isolation

We extracted total DNA from fecal samples and negative (e.g., buffer-only) controls (n = 5) using a solid phase reversible immobilization (SPRI) magnetic bead extraction method (Vo and Jedlicka 2014) with modifications as described in Baiz et al. (2023). Briefly, we aseptically transferred 50mg of fecal material into a 2mL bead-lysis tube containing 0.25g of 0.1mm and 0.25g of 0.5mm zirconia-silica beads. We added 818µL of warmed (65°C) lysis buffer to each bead-lysis tube and homogenized samples using a Precellys 24 Touch (Bertin) for three cycles at 6800rpm for 30s each with a 30s rest time between cycles. We collected and transferred the supernatant into a new 1.5mL tube and incubated samples with Qiagen Solution C3 (Qiagen DNeasy PowerSoil Cat. No. 12888-100-3). After centrifugation, the supernatant was incubated with SPRI beads (1.7x bead-to-supernatant ratio) then washed three times with 80% ethanol. Finally, we eluted the DNA in 10mM Tris-HCl. We quantified the extracted DNA with a Qubit 3.0 Fluorometer (Invitrogen) using the dsDNA HS Assay Kit (Invitrogen, Cat. No. Q32854) and then stored it at -20°C before amplicon sequencing and library preparation.

### 16S Amplicon Sequencing

We performed initial amplification of the V4 region of the bacterial 16S rRNA gene (Caporaso et al. 2012) using 10µM universal primers (515F/806R) with overhanging Illumina adapters (overhang is underlined; 16S-515FB-O: <u>TCGTCGGCAGCGTCAGATGTGTATAAGAGACAG</u>GTGYCAGCMGCCGCGGTAA; 16S-806RB-O: <u>GTCTCGTGGGCTCGGAGATGTGTATAAGAGACAG</u>GGACTACNVGGGTWTCTAAT). Each initial reaction was 25µL, was performed in triplicate, and consisted of 13.5µL nuclease free water, 5µL of 5X Platinum II PCR Buffer (Invitrogen, Cat. No. 14966001), 0.5µL of 10mM dNTP mix, 1.25µL of each primer (10µM each), 0.2µL of Platinum II Taq Hot-Start DNA Polymerase (Invitrogen, Cat. No. 14966001), and 3.3µL of DNA. We also included two negative PCR controls. Reaction conditions were as follows: initial denaturation at 94°C for 2 min; 34 cycles of 98°C for 5s and 68°C for 15s; single final extension at 68°C for 5 min; hold at 12°C. We then pooled the triplicate reactions (total volume = 75µL per sample) and purified them by incubating them with a 1x volume of SPRI beads, followed by two 80% ethanol washes, and finally eluting the bound DNA in 50µL of 10mM Tris-HCl. We ran a 1.5% agarose gel to evaluate amplification and purification.

Following initial amplification, we then performed a second PCR to append Illumina TruSeq i7 and i5 indexes to each pre-cleaned library. Reactions were 30µL and contained 3µL of each 10µM TruSeq primer, 15µL of KAPA HiFi HotStart Ready Mix (Roche, Cat. No. 07958935001), and 9µL of purified DNA from the initial PCR. Reaction conditions were as follows: initial denaturation at 98°C for 45s; 6 cycles of 98°C for 15s, 60°C for 15s, and 72°C for 15s; single final extension at 72°C for 1 min; hold at 12°C. We then purified the indexed PCR product using a double-sided SPRI bead procedure. We first removed potential high molecular weight off-target DNA by incubating indexed PCR products with a 0.75x volume of SPRI beads. We then incubated the supernatant with a 1x volume of SPRI beads to remove potential low molecular weight off-target DNA, followed by two 80% ethanol washes. The final DNA was eluted in 30µL of 10mM Tris-HCl. We ran a 1.5% agarose gel to evaluate amplification and purification, and quantified final products using Qubit as described above.

We normalized library concentrations and pooled all samples to ∼2nM. The final pool was submitted to the University at Buffalo Sequencing and Bioinformatics Core. Sequencing was performed on an Illumina MiSeq platform with the MiSeq Reagent Kit v2 (500 cycles) and 250bp paired-end reads.

### 16S rRNA Amplicon Sequence Processing

A total of 23,750,258 (11,875,129 forward and 11,875,129 reverse) raw Illumina reads across 66 samples (including negatives) were trimmed using the cutadapt plugin (Martin 2011) in QIIME2 version 2024.10 (Bolyen et al. 2019). We then used the DADA2 “denoise-paired” pipeline (Callahan et al. 2016) to pair and filter the trimmed reads with modified parameters (--p-trunc-len-f 228; --p-trunc-len-r 180; --p-trunc-q 10). The quality-filtered reads were denoised into 5,410 amplicon sequence variants (ASVs), which were taxonomically identified using the SILVA reference database (version 138 SSURef NR99 515F/806R; Quast et al. 2013). We then removed ASVs of non-bacterial origin (archaea, eukaryotes, mitochondria, and plant chloroplasts), reducing the dataset to 3,659,515 total reads and 4,845 ASVs. We used the R package “decontam” version 1.22.0 (Davis et al. 2018) to identify and remove possible contaminant ASVs. We used the “prevalence” method with a probability threshold of 0.5 to identify ASVs that were more prevalent in our negative control samples than our biological samples. After removal of those ASVs, 3,538,991 reads and 4,836 ASVs remained. We removed negative control samples from our dataset (which retained 56 biological samples and 4,348 ASVs) before analyzing alpha and beta diversity metrics in QIIME2 and other downstream analyses.

### Statistical Analyses

#### Alpha and Beta Diversity

We discarded 12 samples with fewer than 8,024 reads; this value was determined by a plateau in observed ASV counts beyond this point. The remaining samples were subsampled without replacement to 8,024 reads to generate a “subsampled” ASV table. Then, following Schloss (2024), this process was repeated 100 times (i.e., rarefaction), and for each iteration, we calculated alpha diversity metrics (i.e., within-sample differences; Shannon, Chao1, and Faith’s Phylogenetic Diversity) and beta diversity metrics (i.e., between-sample differences; Bray-Curtis, Jaccard, weighted UniFrac, and unweighted UniFrac). We computed the mean value of each diversity metric across all iterations for each sample, which was used in downstream modeling analyses.

Prior to model building, we tested for multicollinearity among predictors using variance inflation factors and Pearson’s correlation; all values were below common thresholds (GVIF < 5; |r| < 0.7), indicating no problematic correlations. Sampling year was excluded because only one year had associated color data.

We built generalized linear models (GLMs) to evaluate how host color metrics influenced the three different alpha diversity indices. Predictor variables (i.e., fixed effects) included sampling location (state: New York or Pennsylvania), cloacal protuberance score, average bib area, average mask area, and breast feather hue, brightness, and saturation. We used the R package “glmulti” (version 1.0.8; Calcagno and Mazancourt 2010) to generate models with all possible combinations of predictors (n = 128 candidate models per univariate response) and the package “AICcmodavg” (version 2.3-4; Mazerolle 2023) to compare all candidate models to an intercept-only (i.e., null) model using the Akaike Information Criterion corrected for small sample sizes (AICc). Models with ΔAICc scores < 2 were considered equivalent (Burnham and Anderson 2002). For predictors retained in the top models, we examined model-averaged parameter estimates and 85% confidence intervals (CIs); predictors whose 85% CIs did not overlap zero were considered informative (Arnold 2010).

We followed a similar model selection framework modified for multivariate responses to evaluate how host color metrics (same metrics as were used for alpha modelling) influenced the four metrics of beta diversity. Specifically, for each distance matrix, we conducted distance-based redundancy analysis (db-RDA) models for all possible combinations of predictors using the R package “vegan” version 2.6-10 (Oksanen et al. 2025) and *capscale* function. For each model, we calculated the adjusted R^2^, conducted a global permutation test (999 permutations), and performed marginal tests of individual predictors. We retained only the top 10% of models ranked by adjusted R^2^ to focus on the models with the highest explanatory value and reduce noise. We then summarized predictor importance across these top models by quantifying their frequency of occurrence, average marginal effects, and representation among models that had a significant p-value from the permutation test. Alpha and beta analyses were performed with (1) New York and Pennsylvania samples combined and (2) New York samples only, due to a larger New York sample size, to assess the robustness of the results.

#### Differential Abundance and Microbial Associations

We used Microbiome Multivariable Associations with Linear Models (MaAsLin2 version 1.16; Mallick et al. 2021) to identify ASVs that were differentially abundant among and/or associated with host plumage features, including average bib area, average mask area, and breast feather hue, brightness, and saturation. We included sampling location and cloacal protuberance score as additional fixed effects. As input for our models, we generated 100 “subsampled” ASV tables, each subsampled to 8,024 reads (rarefaction; described above and in Schloss [2024]). For each ASV in each sample, we calculated the mean abundance across all 100 tables, producing a single “mean ASV table” that was used for these analyses (retaining 4,088 ASVs). We converted our mean ASV table to relative abundances, and then specified the CPLM (Compound Poisson [generalized] Linear Model) analysis method because our data were zero-inflated (i.e., most features were rare). We assigned the minimum abundance threshold to 0.0001 and the minimum prevalence threshold to 0.15 (all samples) or 0.20 (NY samples only). We specified “none” for normalization because relative abundance data were used as input and “none” for transformation because of the intrinsic log link of CPLM. Continuous fixed effects were Z-scored for standardization. We applied the Benjamini-Hochberg correction for multiple hypothesis testing using a false discovery rate (FDR) threshold of 0.1 and 0.05 (i.e., Q-values ≤ 0.1 and 0.05 were considered significant or highly significant, respectively).

## RESULTS

### Phenotypic Diversity

Our sampling captured a wide range of phenotypes (Table 1; Figure S1), in both color and size, consistent with the range of variation for these traits documented in previous studies (Tarof et al. 2005, Dunn et al. 2008, 2010).

**Table 1.** Summary of colorimetric and phenotypic traits of all male Common Yellowthroat samples (New York and Pennsylvania), including the mean, standard deviation (SD), and standard error (SE).

|  | Mean | SD | SE |
| --- | --- | --- | --- |
| Hue | 0.743 | 0.053 | 0.007 |
| Brightness | 0.17 | 0.044 | 0.006 |
| Saturation | 0.737 | 0.023 | 0.003 |
| Average bib area (cm<sup>2</sup>) | 11.763 | 2.968 | 0.396 |
| Average mask area (cm<sup>2</sup>) | 2.363 | 0.442 | 0.059 |

### Taxonomic Diversity

Our mean ASV table contained 4,088 ASVs from 37 bacterial phyla (including the category “Unassigned”). The most common phylum was Proteobacteria, which represented 56.7% of the total reads across samples, followed by Actinobacteriota (18.9%) and Firmicutes (15.7%; Figure 1). The most common family was Enterobacteriaceae (phylum Proteobacteria) and represented 12.6% of total reads. The two most prevalent ASVs were in the families Pseudonocardiaceae and Microbacteriaceae (both in the phylum Actinobacteriota), which were found in 31 out of 44 samples (70.5%).

**Figure 1.**
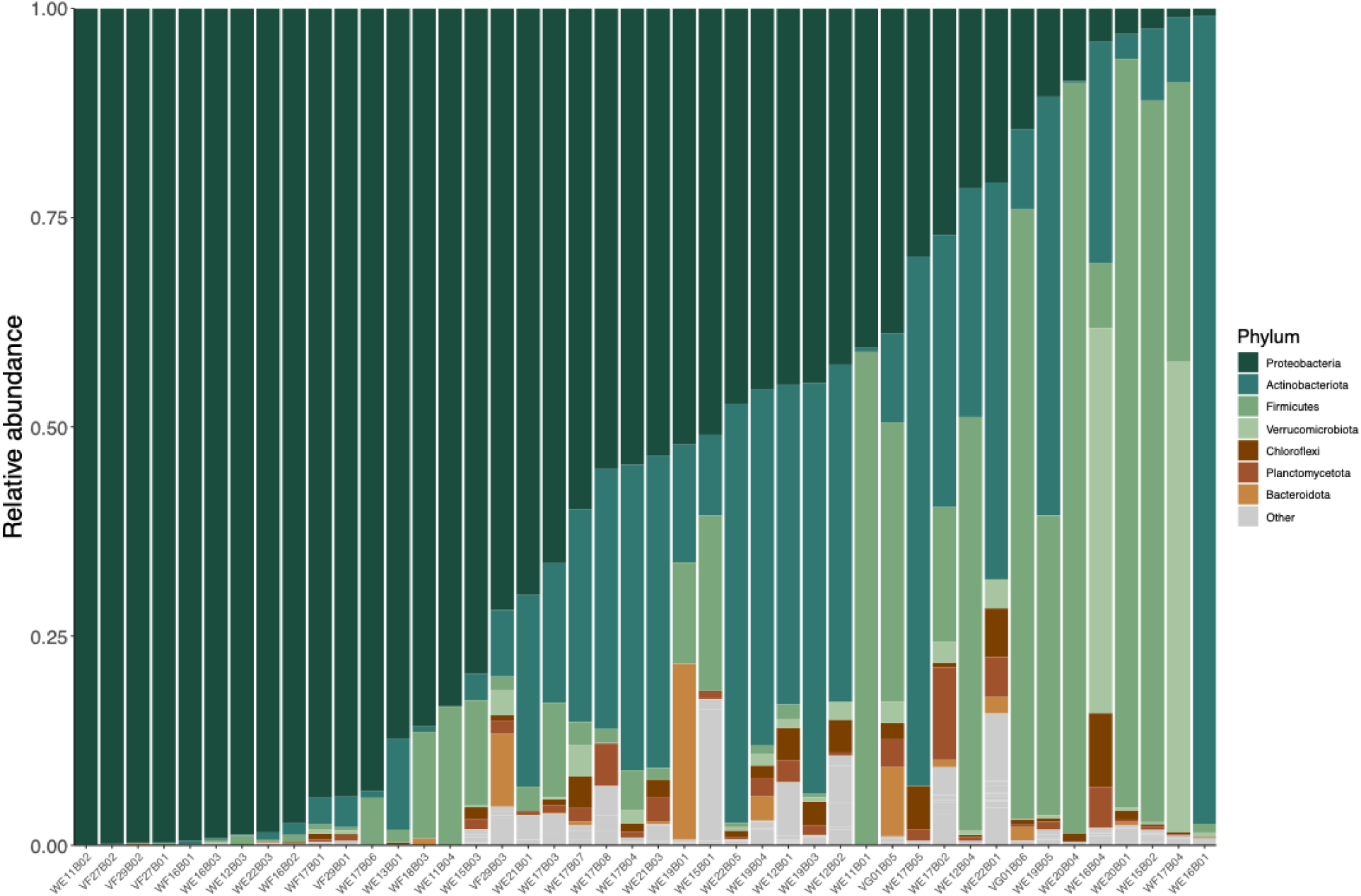
Stacked barplots depicting the relative abundance of bacterial phyla in the rarefied dataset. Only the seven most abundant phyla are displayed; all remaining taxa are grouped as “Other.” Each column represents an individual sample and is ordered by decreasing relative abundance of the most abundant phylum (Proteobacteria).

### Alpha Diversity

For Shannon diversity, analyses using all samples yielded 14 competitive models (ΔAICc ≤ 2), none of which were the intercept-only model (ΔAICc = 41.2). Across the competitive models (Table 2), cloacal protuberance score was the most consistently supported predictor, appearing in 12/14 (86%) of competitive models (Figure 2A). Its 85% CIs excluded zero for a score of “3” in all models and for a score of “2” in 7/12 (58%) of models. In contrast, the New York-only subset produced 10 competitive models, including the intercept-only model (ΔAICc = 0.6), indicating that none of the predictors substantially improved model fit when Pennsylvania samples were removed (Table 3).

**Figure 2.**
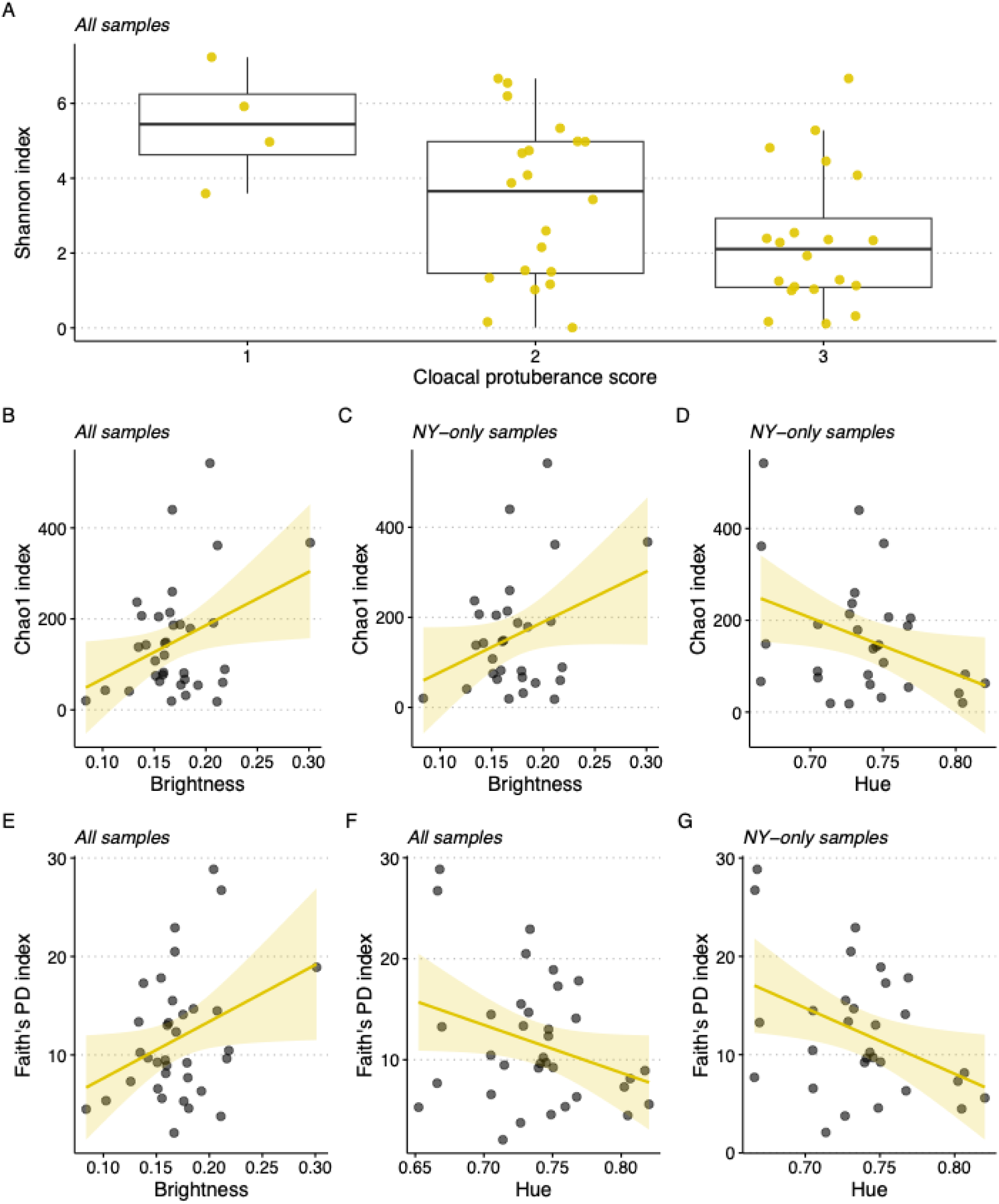
Plots illustrating the relationships between Common Yellowthroat gut microbial diversity indices and the most informative predictors (e.g., present in at least 50% of top models and were significant in at least 50% of those top models) identified from the model selection process. Plots are organized by microbiome alpha metric response (Shannon [A], Chao1 [B-D], and Faith’s PD [E-G]). The specific dataset (either all samples or New York-only samples) is indicated in plot subtitles. Plots display raw data (with 95% confidence intervals in scatterplots), not fitted models, for simplicity; thus, no statistical metrics are denoted within the plots.

**Table 2.** Summary of top models (ΔAICc ≤ 2) for alpha diversity using all samples (New York and Pennsylvania). Each alpha diversity metric lists the predictors in the top models, their frequency among the top models, the proportion of those models in which the 85% confidence intervals (CIs) for the predictor’s estimate excluded zero, and the direction of the effect for predictors with at least one 85% CI excluding zero. Predictors with no 85% CIs excluding zero are indicated with “NA” in the Effect Direction column. Predictors not included in any top models for a given metric are not listed.

| **Metric** | **Predictor** | **Frequency in top models** | **% of top models that 85% CIs exclude zero** | **Effect Direction** |
| --- | --- | --- | --- | --- |
| Shannon | cp (3) | 12/14 (86%) | 12/12 (100%) | Negative |
|  | cp (2) | 12/14 (86%) | 7/12 (58%) | Negative |
|  | hue | 7/14 (50%) | 0/7 (0%) | NA |
|  | average mask area | 6/14 (43%) | 5/6 (83%) | Positive |
|  | state (PA) | 5/14 (36%) | 2/5 (40%) | Negative |
|  | saturation | 4/14 (29%) | 0/4 (0%) | NA |
|  | average bib area | 3/14 (21%) | 1/3 (33%) | Positive |
|  | brightness | 3/14 (21%) | 0/3 (0%) | NA |
| Chao1 | brightness | 4/5 (80%) | 4/4 (100%) | Positive |
|  | hue | 2/5 (40%) | 1/2 (50%) | Negative |
|  | average mask area | 1/5 (20%) | 0/1 (0%) | NA |
|  | saturation | 1/5 (20%) | 0/1 (0%) | NA |
| Faith's PD | brightness | 5/8 (63%) | 5/5 (100%) | Positive |
|  | hue | 4/8 (50%) | 3/4 (75%) | Negative |
|  | state (PA) | 2/8 (25%) | 0/2 (0%) | NA |
|  | saturation | 1/8 (13%) | 0/1 (0%) | NA |
|  | average mask area | 1/8 (13%) | 0/1 (0%) | NA |
|  | average bib area | 1/8 (13%) | 0/1 (0%) | NA |

**Table 3.** Summary of top models (ΔAICc ≤ 2) for alpha diversity using only New York samples. Each alpha diversity metric lists the predictors in the top models, their frequency among the top models, the proportion of those models in which the 85% confidence intervals (CIs) for the predictor’s estimate excluded zero, and the direction of the effect for predictors with at least one 85% CI excluding zero. Predictors with no 85% CIs excluding zero are indicated with “NA” in the Effect Direction column. For metrics where the intercept-only model was among the top models (e.g., Shannon), no individual predictors were included in top models, so all entries are shown as “NA.” Predictors not included in any top models for a given metric are not listed.

| **Metric** | **Predictor** | **Frequency in**<br>**top models** | **% of top models that**<br>**85% CIs exclude zero** | **Effect**<br>**Direction** |
| --- | --- | --- | --- | --- |
| Shannon | NA | NA | NA | NA |
| Chao1 | hue | 3/5 (60%) | 3/3 (100%) | Negative |
|  | brightness | 3/5 (60%) | 2/3 (66%) | Positive |
|  | saturation | 1/5 (20%) | 1/1 (100%) | Positive |
|  | average bib area | 1/5 (20%) | 0/1 (0%) | NA |
| Faith's PD | hue | 3/3 (100%) | 3/3 (100%) | Negative |
|  | average bib area | 1/3 (33%) | 0/1 (0%) | NA |
|  | brightness | 1/3 (33%) | 0/1 (0%) | NA |

For Chao1 diversity analyses using all samples yielded five competitive models, which were all strongly supported relative to the intercept-only model (ΔAICc = 114.4). Breast feather brightness appeared in 4/5 (80%) of competitive models, and its 85% CIs did not overlap zero in any, suggesting a consistent positive effect of brightness on Chao1 diversity (Table 2; Figure 2B). With the New York-only subset, there were five competitive models, which were all supported relative to the intercept-only model (ΔAICc = 2.3). Breast feather brightness (Figure 2C) and hue (Figure 2D) both appeared in 3/5 (60%) of competitive models; the 85% CIs of brightness did not overlap zero in 2/3 (67%) of models and the 85% CIs of hue did not overlap zero in any of those models, suggesting a regularly consistent positive effect of brightness and negative effect of hue on Chao1 diversity (Table 3).

For Faith’s PD, analyses using all samples yielded eight competitive models, which were all strongly supported relative to the intercept-only model (ΔAICc = 63.8). Breast feather brightness appeared in 5/8 (63%) of competitive models, and its 85% CIs did not overlap zero in any, suggesting a consistent positive effect of brightness on Faith’s PD (Table 2; Figure 2E). Hue also appeared in 4/8 (50%) of competitive models with its 85% CIs not overlapping zero in 3/4 (75%) of models, suggesting a consistent negative effect (Table 2; Figure 2F). With the New York-only subset, there were three competitive models, which were all supported relative to the intercept-only model (ΔAICc = 2.9). Breast feather hue appeared in all competitive models and its 85% CIs did not overlap zero in any models, suggesting a consistent negative effect of hue on Faith’s PD (Table 3, Figure 2G). Full model results can be found in Table S1.

### Beta Diversity

Beta diversity analyses based on Bray-Curtis dissimilarity showed limited support for any predictors. In the full dataset (all samples), the seven models comprising the top 10% of adjusted R² values were all non-significant in global tests (p > 0.05). However, breast feather hue appeared consistently across these models despite having non-significant marginal effects (Table 4). A similar pattern was observed in the New York-only dataset: the seven top models showed no global significance, and breast feather saturation, hue, and average mask area were the most frequently retained predictors, though none exhibited significant marginal effects (Table 5).

**Table 4.**
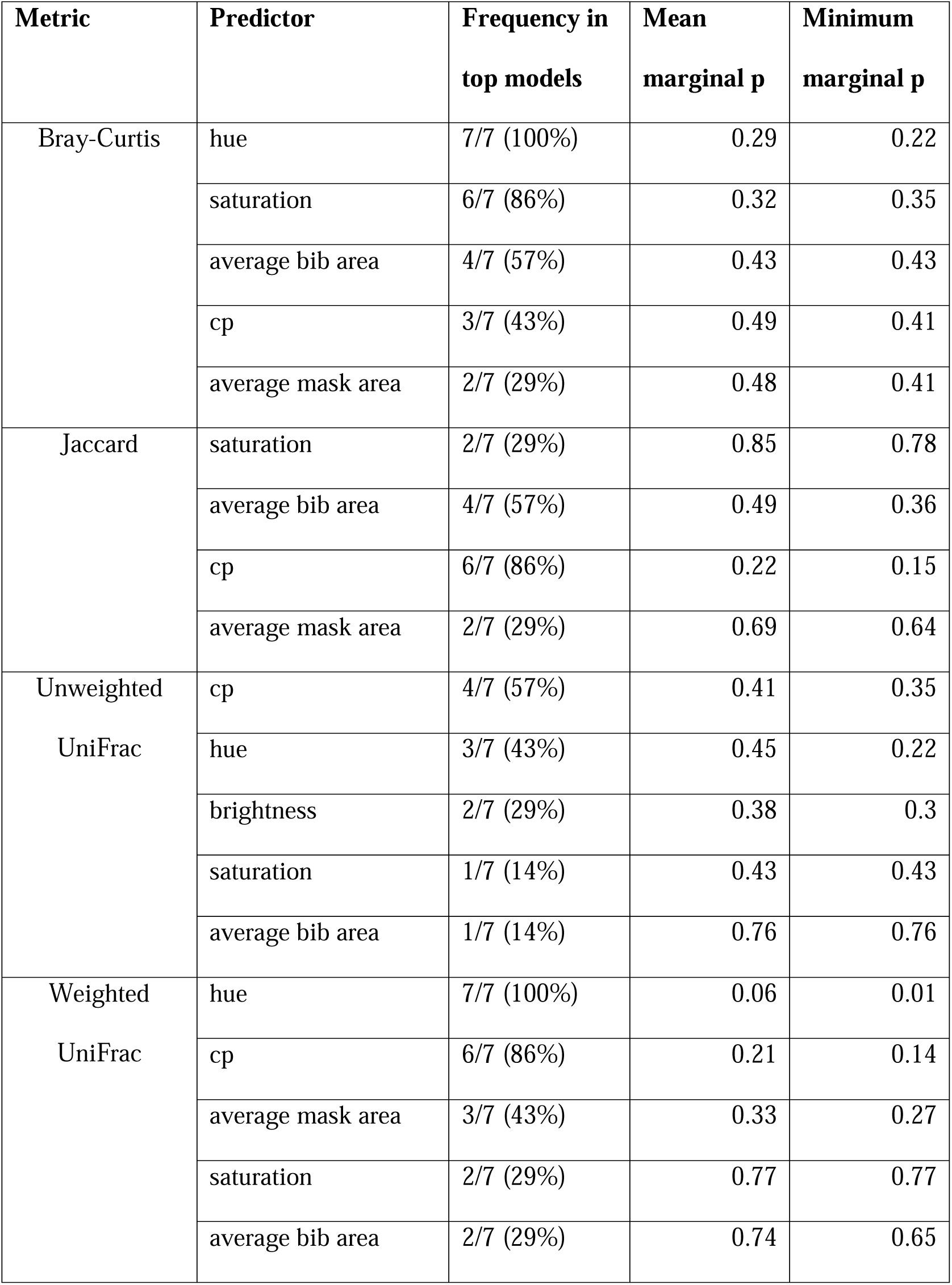
Summary of top models (top 10% of adjusted R² values) for beta diversity using all samples (New York and Pennsylvania). Each beta diversity metric lists the predictors included in the top models, the frequency of occurrence among the top models, the mean marginal p-value across top models, as well as the minimum marginal p-value across top models. Predictors not included in any top models for a given metric are not listed.

**Table 5.**
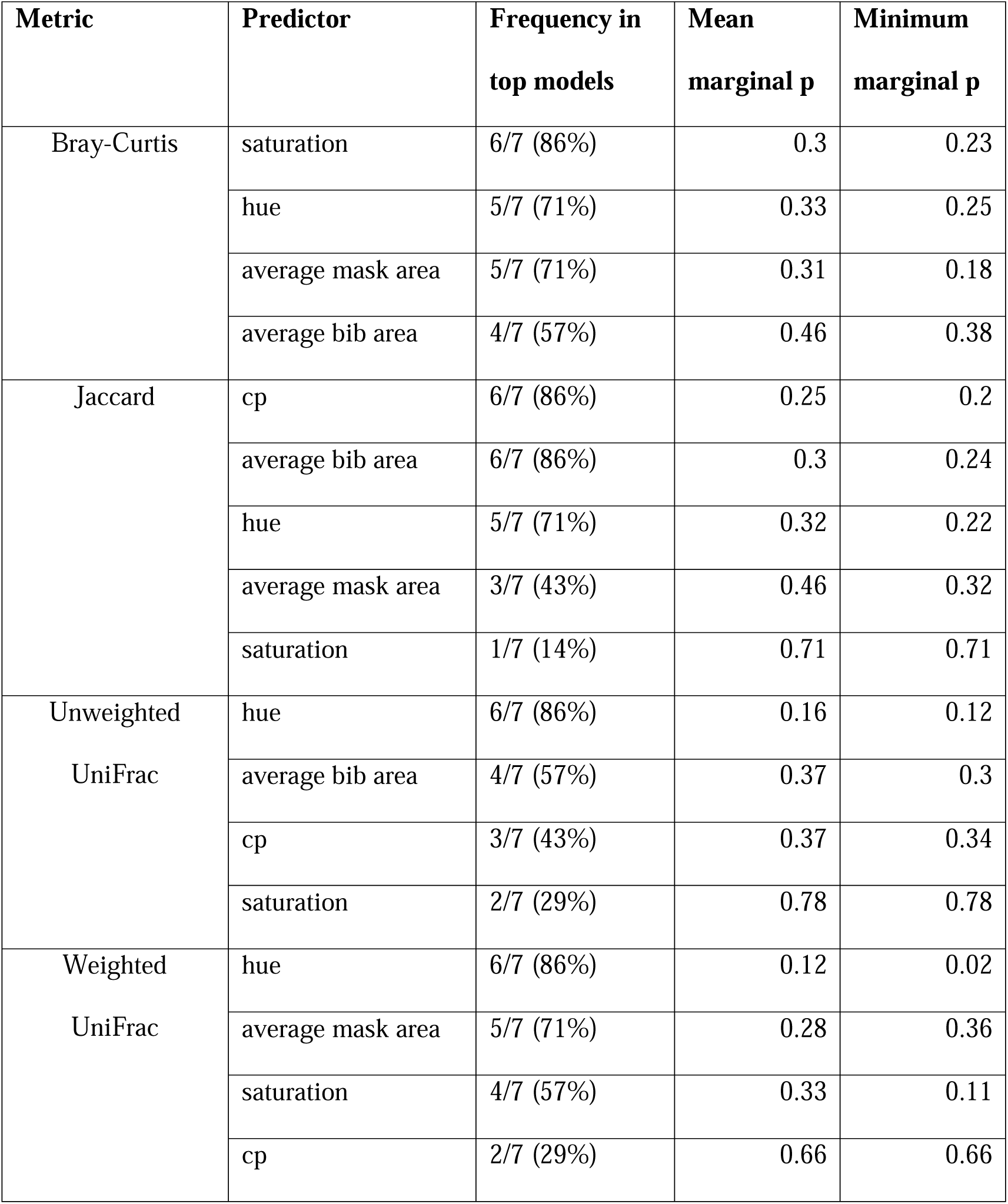
Summary of top models (top 10% of adjusted R² values) for beta diversity using only New York samples. Each beta diversity metric lists the predictors included in the top models, the frequency of occurrence among the top models, the mean marginal p-value across top models, as well as the minimum marginal p-value across top models. Predictors not included in any top models for a given metric are not listed.

Jaccard-based beta diversity showed similar patterns of limited explanatory power. In the full dataset, the seven models comprising the top 10% of adjusted R² values were all non-significant in global tests. Cloacal protuberance score appeared frequently across these models (6/7; 86%) but exhibited no significant marginal effects (Table 4). Similarly, in the New York-only dataset, none of the seven top models were globally significant, and average mask area and cloacal protuberance score were the most common predictors (both in 6/7 [86%] of models). However, neither showed significant marginal effects (Table 5).

Unweighted UniFrac results also indicated weak predictor support. Across all samples, none of the seven top models comprising the top 10% of adjusted R² values were globally significant, and the most frequent predictor (cloacal protuberance score [4/7; 57%]) did not exhibit significant marginal effects (Table 4). In the New York-only subset, the seven top models were also not globally significant; breast feather hue appeared most frequently (6/7; 86%), but its marginal effects were non-significant (Table 5).

Weighted UniFrac analyses revealed somewhat stronger patterns than the other beta diversity metrics. In the full dataset, three of the seven top models were globally significant, with breast feather hue in all seven models (including all three significant models) and cloacal protuberance score appearing in six (including two significant models). For hue, marginal effects were consistent and significant or approaching significance across all seven models (mean marginal p = 0.06; minimum = 0.01; Table 4) and were significant in all three globally significant models (mean marginal p = 0.03; minimum = 0.01; Table S2), suggesting a possible influence of hue on community structure. Cloacal protuberance score showed weaker support across models (Table 4). In the New York-only dataset, four of the seven top models were globally significant. Hue was retained in six of these models (including three significant models), and average mask area was retained in five (including two significant models). Hue continued to exert some influence on community structure in the New York-only data as its marginal effects were significant or approaching significance across all seven models (mean marginal p = 0.12; minimum = 0.02; Table 5) and were significant in two of three globally significant models (mean marginal p = 0.05; minimum = 0.02; Table S2). Average mask area showed weaker support in this dataset (Table 5). Full model results can be found in Table S2.

### Differential Abundance and Microbial Associations

Using the full dataset (i.e., New York and Pennsylvania samples combined), differential abundance analyses in MaAsLin2 identified 22 ASVs whose abundances were significantly associated with various host plumage features (Q < 0.1), of which eight were significant at the alpha level of 0.05 (Table S3). These ASVs were classified into 10 (Q < 0.1) and six (Q < 0.05) genera. Among these genera, *Methylobacterium-Methylorubrum* had the greatest number of significant associations across all predictors (Figure 3). Of those significant associations, mask area (positive), breast feather hue (negative), and saturation (negative) had the strongest effects (Figure 4). Mask area also has a strong positive association with *Rickettsia* (Figure 3).

**Figure 3.**
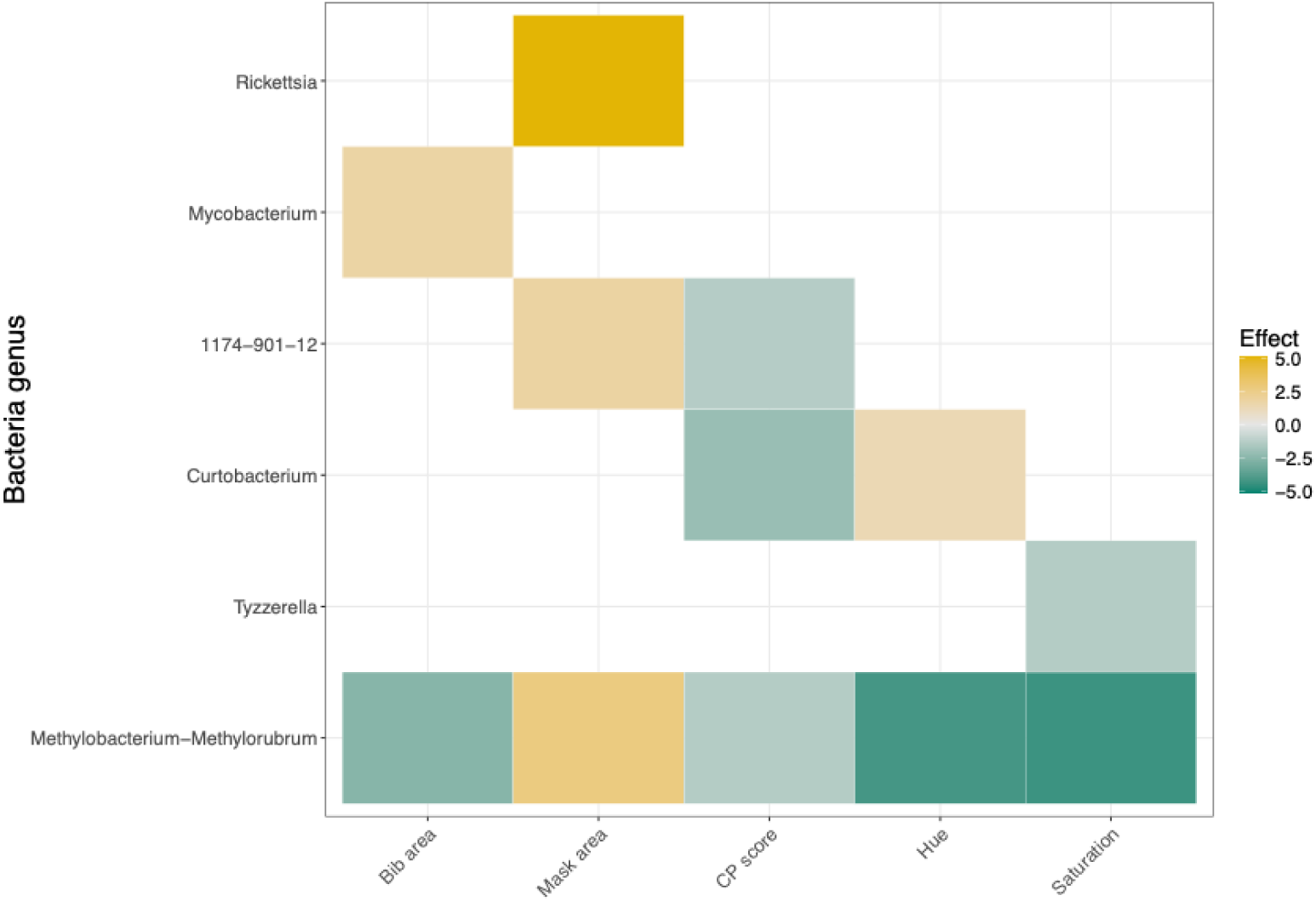
Heatmap illustrating ASV genera that are most strongly (Q < 0.05) associated with host plumage features. Colors represent the strength and direction of the association (effect score = −log10(Q-value) × sign(coefficient)). Teal represents negative associations and gold represents positive associations.

**Figure 4.**
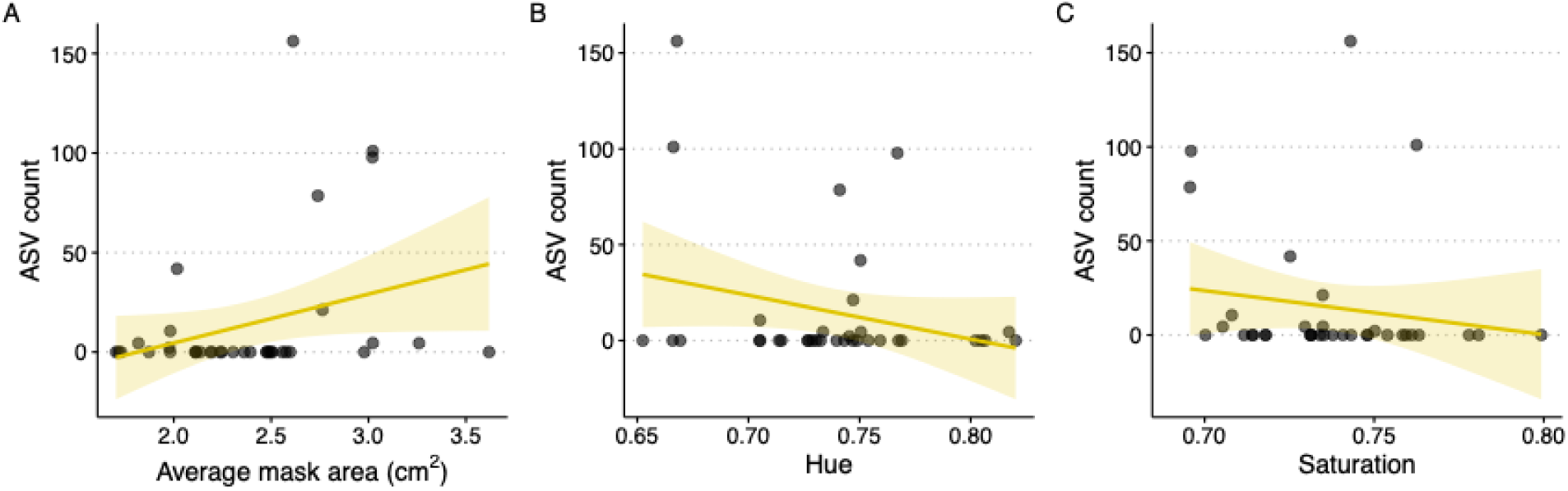
Counts of *Methylobacterium-Methylorubrum* (sum of congeneric ASV read counts) in relation to three host plumage traits that were identified as strongly associated with this genus(average mask area [A], feather hue [B], and feather saturation [C]). Each point represents a sample and trend lines represent linear regression fits with 95% confidence intervals.

Using the New York-only dataset, three ASVs were significantly associated with three different host plumage traits (Q < 0.1; Table S3). These three ASVs represented three different genera (Figure S2A), of which two were significant at Q < 0.05 (*Rickettsia* and *Kineococcus*). *Rickettsia* had the strongest association, a positive correlation with mask area, but this was driven by a single sample (Figure S2B).

## DISCUSSION

Across individuals, we observed substantial variation in both gut microbiome diversity and host phenotypes. The overall gut microbiome profiles in our study mirror findings in previous studies of breeding warblers (Skeen et al. 2021, Baiz et al. 2023, 2024, Trevelline et al. 2023), and other passerines (Skeen et al. 2023, Slevin et al. 2025). Specifically, Proteobacteria was the most abundant phylum, followed by Actinobacteria and Firmicutes. Despite these broad-level similarities, the identity of the dominant phylum varied widely among individuals (Figure 1), which is consistent with high levels of inter-individual variation reported in populations of wild birds (Bodawatta et al. 2021, Baiz et al. 2023, Liukkonen et al. 2024). Similarly, we observed substantial variation across all measured phenotypic traits, including breast feather colorimetric traits, cloacal protuberance score, and mask and bib area (Figure S1). In many cases, values spanned two- to three-fold differences between individuals (Table 1). The degree of variability is characteristic of ornaments in this species and aligns with previously documented patterns of phenotypic diversity (Dunn et al. 2008, 2010, Sly et al. 2022). Thus, the extensive individual microbial and phenotypic variation underscores the dynamic nature of these traits and provides valuable context for interpreting the associations we detected between them.

We found that plumage coloration consistently emerged as an important correlate of gut microbiome diversity (e.g., was present in at least 50% of top models and significant in at least 50% of those top models). Specifically, breast feather hue and brightness were the most frequently retained predictors in the top models for alpha diversity metrics in both the full dataset and New York-only dataset, including Chao1 (ASV richness) and Faith’s Phylogenetic Diversity (sum of branch lengths in the phylogeny of all ASVs), and were also almost always significant when retained (Table 2, Table 3). By contrast, the few other studies testing relationships between colorimetric plumage traits and microbiome variation have found no significant associations with breast feather color metrics; however, these studies focused on Northern Cardinals (*Cardinalis cardinalis*; Slevin et al. 2025) and Common Waxbills (*Estrilda astrild*; Marques Silva et al.2023), which both have red breast feathers (as opposed to yellow in Common Yellowthroats). Red and yellow feather pigmentations are mediated by different biochemical and genetic processes (e.g., yellow carotenoids like β-carotene are converted to red ketocarotenoids like canthaxanthin via a ketolase), which may contribute to these differences (Mundy et al. 2016, Lopes et al. 2016, Toews et al. 2017). Most beta metric models yielded non-significant marginal effects of predictors, suggesting that their influence on among-individual variation may be weak or difficult to detect with the present data. The only exception was hue, which was frequently significantly associated with Weighted UniFrac diversity (a phylogenetically informed metric that weights branch lengths by ASV abundance; Table 4). Taken together, these patterns suggest that color traits are linked not only to the variation in gut microbiome richness, but also to its phylogenetic structure.

Notably, these associations were specific to colorimetric traits rather than ornament size for either carotenoid- (bib) or melanin-based (mask) traits. Overall, male Common Yellowthroats with brighter (higher brightness values) and more orange-tinted (lower hue values) breast feathers harbored richer and more phylogenetically diverse gut microbiomes, whereas bib and mask size showed little evidence of association with microbiome variation. Experimental evidence in Common Yellowthroats demonstrated that ornament size (both mask and bib) was more important than coloration for both inter- and intrasexual selection (Tarof et al. 2005, Dunn et al. 2008). Thus, our results suggest that gut microbiomes may be more closely tied to physiological processes underlying carotenoid pigment acquisition (e.g., from external sources) and processing (e.g., absorption, transport, modulation, deposition) rather than to ornament size, the stronger target of sexual selection in this species.

Although there are several potential mechanisms that could explain this association, it is important to note that the directionality of the relationship remains unclear. One possibility is that variation in the gut microbiome can influence a host’s ability to acquire, metabolize, or regulate carotenoids through effects on digestion, gut chemistry, and nutrient processing. Because carotenoids are absorbed in the digestive tract, gut microbial activity could shape pigment bioavailability directly by altering absorption efficiency (Grolier et al. 1998, Djuric et al. 2018), indirectly by modifying the gut metabolic environment (Karlsson et al. 2012, Pudlo et al. 2015), or through a combination of direct and indirect factors (Bohn et al. 2017). An alternate, but related, explanation is that microbiome diversity and carotenoid-based coloration covary because they are influenced by the same underlying ecological and physiological factors (e.g., environmental health, diet quality, hormones, immunity). For example, ecological stressors such as urbanization can reduce the concentration of naturally available carotenoids (Isaksson and Andersson 2007) while also impacting gut microbiome diversity (Phillips et al. 2018). Similar patterns have been observed in hosts experiencing physiological stress (Dunn et al. 2010, Knutie 2020, Bernabeu et al. 2024), suggesting that microbiome diversity and carotenoid-based coloration may respond in parallel to shared physiological pressures (Slevin et al. 2026). Taken together, our results suggest that these traits are tightly physically and functionally linked, likely through interactions mediated by the environment and host physiology, even if the causal mechanisms remain unresolved.

Another possible mechanism linking microbiome diversity and variation in plumage coloration could be related to host behavior and the transmission of microbes. For example, brighter or more orange-tinted males may engage in more frequent social, territorial, or reproductive interactions (including extra-pair copulations), which increases transmission opportunities for microbes among individuals and could ultimately inflate gut microbiome diversity, particularly during the breeding season, which is consistent with the patterns we observed. Such socially mediated microbial exchanges have been implicated in observational (Levin et al. 2016, Escallón et al. 2019) and experimental studies (Taff et al. 2021). In Rufous-Collared Sparrows (*Zonotrichia capensis*), microbiome diversity increases during the breeding season and is followed by a reduction at its conclusion (Escallón et al. 2019). This cyclical pattern in diversity has been attributed to extra-pair matings and/or to hormonal (and ultimately, behavioral) changes in this species (Escallón et al. 2017). In Common Yellowthroats, extra-pair matings are quite common (∼50%) and have been shown to be correlated with mask size (Thusius et al. 2001a, b) and bib brightness (Freeman-Gallant et al. 2010). Thus, it is possible that microbial transmission could occur during mating or social interactions and provide a potential behavioral route by which plumage coloration covaries with gut microbiome diversity, though subsequent (and ideally manipulative) work is required to further test this possibility. Beyond coloration, our results indicate that other aspects of reproductive physiology may also shape gut microbiome diversity.

Cloacal protuberance (CP) score (i.e., height) was consistently and significantly negatively associated with gut microbiome Shannon diversity (richness and evenness) when we analyzed the full dataset (Figure 2). The presence of a CP is indicative of breeding condition status in male birds (and circulating testosterone levels; Biswas et al. 2007), where swelling is absent during most of the annual cycle (pre- and post-breeding) but develops and enlarges upon entering breeding condition to serve a role in sperm storage and assist in copulation, then regresses (Pyle 2022). Our sampling ranged from mid-May through early July, thus we captured the entire breeding season as well as the complete range of reproductive states via CP scoring. Notably, we recorded all three CP scores (low, intermediate, and high height) within a single week in mid to late-May. This pattern suggests that the significant relationship between CP score and microbiome diversity cannot be attributed solely to sampling date, but instead reflects meaningful biological differences among individuals transitioning through reproductive states.

There are several non-mutually exclusive mechanisms that could explain the decline in microbiome diversity as individuals progressed through reproductive states. One possibility involves physiological changes associated with the annual cycle. Variation in avian gut microbiome diversity across the annual cycle has been observed in several other studies (Escallón et al. 2019, Skeen et al. 2021, 2023, Trevelline et al. 2023), particularly during the transitions between migration, breeding, and non-breeding periods (reviewed in Capilla-Lasheras and Risely 2025). In a longitudinal study of Kirtland’s Warblers (*Setophaga kirtlandii*), individuals sampled prior to spring migration on the non-breeding grounds (March and April), shortly after arrival on the breeding grounds (May and June), and later in the breeding season (July) exhibited a progressive decline in alpha diversity, suggesting physiological remodeling of the gut microbiome as birds transition from migratory to reproductive states (Skeen et al. 2021). Although our sampling was restricted to the breeding season and represents independent observations rather than repeated measures, the similar decline observed here aligns with evidence that microbiome communities are restructured as hosts shift into breeding condition. This shift may be associated with increased energetic investment in mate acquisition, territorial defense, and parental care and could potentially favor a narrower subset of microbial taxa. However, future studies that integrate metagenomic or metabolomic approaches will be essential to determine if the compositional shifts observed here during the early breeding season are accompanied by functional shifts, as has been observed during the stages of fall migration in warblers (Trevelline et al. 2023).

Another non-mutually exclusive explanation involves the rapid environmental and dietary changes birds experience upon arrival to breeding grounds following migration. Lewis et al. (2017) demonstrated that the gut microbiome of thrushes sampled at a migratory stopover site quickly converged with one another following their arrival, consistent with the idea that the local environmental conditions and resource use can strongly alter microbial communities. In our study, individuals captured with low CP scores likely represent birds that had only recently arrived at the breeding grounds (mid-May, prior to the onset of active reproduction; Guzy and Ritchison 2020). These individuals may have acquired a diversity of microbiota during migration that subsequently stabilized after settlement on the breeding grounds. This interpretation is corroborated by Skeen et al. (2021), in which microbiome diversity in Kirtland’s Warblers rapidly declined over the first few days following their arrival to the breeding grounds (though it rebounded after approximately one week). Such patterns are supported by the broad geographic range of Common Yellowthroats throughout their annual cycle; populations from our study region likely migrate along the Atlantic Seaboard and may overwinter as far as the Yucatán Peninsula and Jamaica (Bobowski et al. 2025). Exposure to diverse habitats and microbes along this migratory route could contribute to elevated microbiome diversity upon arrival, followed by a reduction as birds transition to more converged diets and local environments on the breeding grounds. Longitudinal sampling and/or an integrative study on both gut microbial and dietary data (Baiz et al. 2023, Uehling and Houtz 2025) may shed light on this possibility.

Certain microbial taxa may directly link the gut microbiome to host coloration. Some gut-associated microbes are capable of synthesizing carotenoid compounds themselves (Eilers et al. 2024), raising the possibility that microbiomes may contribute supplemental pigment sources beyond those obtained through the diet, as has been proposed in several recent studies across a wide array of taxa (Liu et al. 2020, Baiz et al. 2024, Koellsch et al. 2024). Although we did not uncover many microbial taxa whose abundance strongly differed across plumage traits or CP score in either of our datasets, *Methylobacterium-Methylorubrum* was among the most common (Figure 4, Figure 3). *Methylobacterium-Methylorubrum* is a facultative methylotrophic bacterium and a known carotenoid producer (Downs and Harrison 1974, Van Dien et al. 2003). One of best studied representatives of this genus, *Methylobacterium extorquens* AM1, is pink-pigmented and produces a C30 carotenoid (Mo et al. 2023). While the majority of carotenoid pigments extracted from feathers are C40-based (Toomey et al. 2022), the relative abundance of this same genus was positively associated with host plumage yellowness in *Vermivora* warblers (Baiz et al. 2024). Thus, together with our results, this may suggest a direct microbial contribution to pigment availability for hosts. However, the limited number of significantly differentially abundant taxa found herein suggests that microbiome-color relationships do not stem from a single taxon. Instead, the observed relationships may reflect shifts in groups of functionally similar taxa rather than changes in individual ASVs. Such functional redundancy goes undetected using differential abundance analyses based on ASV taxonomy, but functional metagenomic analyses would help to capture this potential signal in the future.

In conclusion, our data indicate that Common Yellowthroat microbiome diversity varies with carotenoid-based plumage coloration traits, namely breast feather brightness and hue, as well as breeding condition. Interestingly, we found no relationships with ornament (i.e., bib, mask) size, the primary target of sexual selection in this species (Thusius et al. 2001b, Tarof et al. 2005, Dunn et al. 2008). Although more work is needed to rule out sexual selection as a primary driver of microbiome variation in this species, our results appear to instead reflect strong physiological interactions between plumage pigmentation and microbes in the gut. Host behavior or environmental conditions may be further contributors to these patterns, either interactively with or independent of physiology. Additional experimental or strategic observational studies could help disentangle cause and effect and clarify the directionality of the relationships we observed. Overall, this study demonstrates that the gut microbiome may represent a key link between internal physiology and external signals that mediate avian sexual selection. Integrative approaches that bridge microbial ecology, physiology, and behavioral biology will be essential to resolving these relationships in wild populations.

## DATA AVAILABILITY

Raw sequence data were submitted to the Sequence Read Archive at the National Center for Biotechnology Information (BioProject PRJNA1379636). Reviewer link: https://dataview.ncbi.nlm.nih.gov/object/PRJNA1379636?reviewer=nm2lbo2sgkftdhc0a6igr3u3l3

## ETHICS STATEMENT

Birds were captured and handled under the United States Geologic Survey Bird Banding Laboratory Permit #24043 and a protocol approved by IACUC at Penn State University (protocol no. 201900879).

## ACKNOWLEDGEMENTS

We thank Nicholas Sly, Peter Dunn, and David Toews for their advice about study design. We also thank Matthew Medler and Derek Rogers for their help in identifying field locations, as well as the New York State Parks and Department of Environmental Conservation, the Nature Conservancy, the Adirondack Land Trust, and Paul Smith’s College for permission to work on their land and preserves. Data were gathered on the unceded traditional territories in Pennsylvania of the Shawnee Tribe in regions now known as central Pennsylvania and in the Lenape Delaware tribe in eastern Pennsylvania. Data were also collected in the present-day Saint Regis Mohawk Tribe territories in and around the Adirondack region of New York State as well as the Oneida territories in the Albany region of New York State. We thank the present-day Indigenous community members living in these regions as well as the land for these data. This work would not have been possible without traveling through and collecting on this tribal land.

## AUTHOR CONTRIBUTIONS

M.D.B. conceived the study and acquired funding. M.D.B., M.M., and L.P. conducted fieldwork to collect samples. A.E.M., M.G.P., and S.G. conducted laboratory work. A.E.M. led data analysis with input from M.D.B. and D.T.B. A.E.M and M.D.B. led manuscript writing. All authors contributed to revisions and approved the final version of the manuscript.

