## Supplementary figures and images for "Colorful connections: pigment-based plumage and breeding condition are associated with gut microbiome variation in the Common Yellowthroat"

### Figure S1

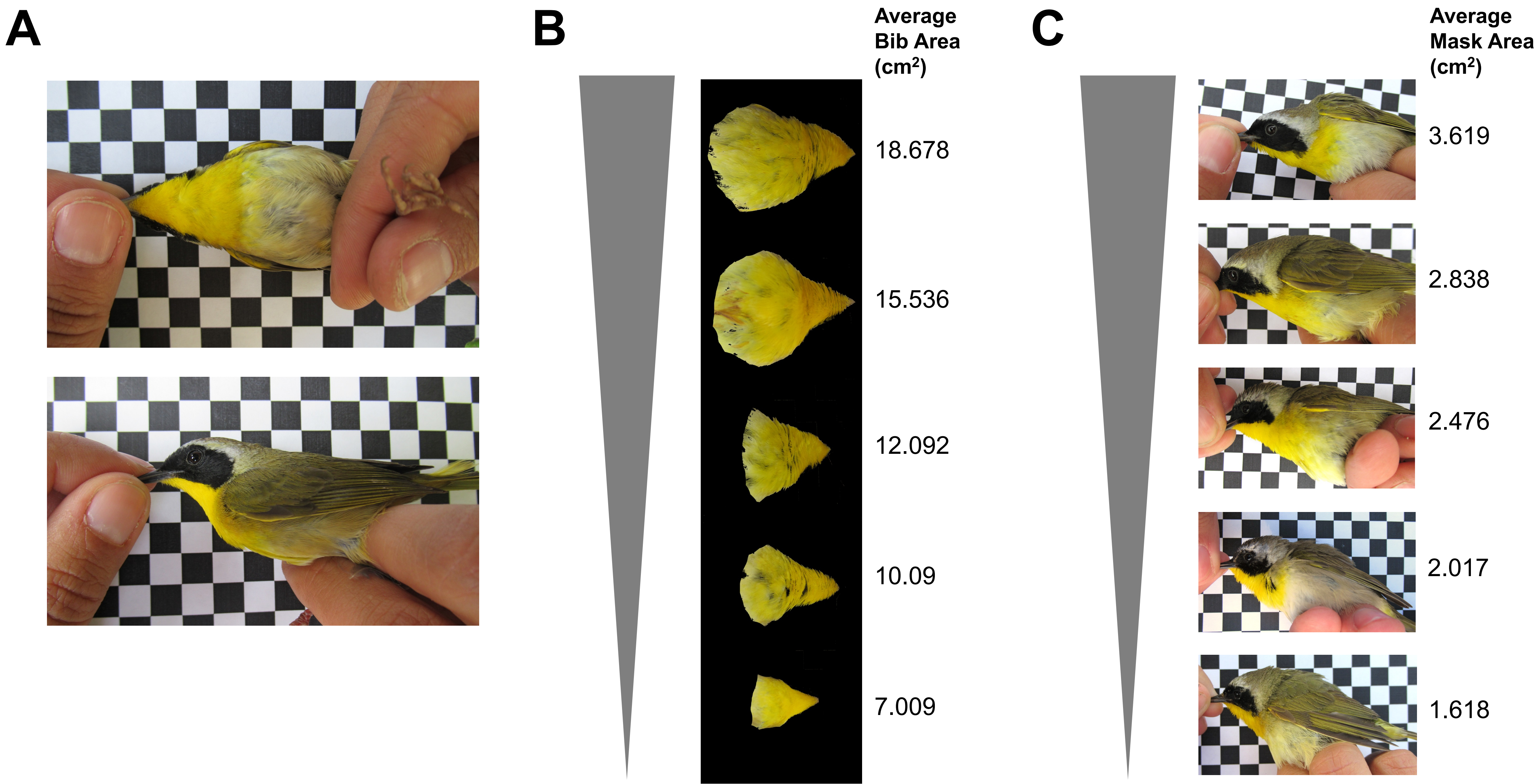

### Figure S2

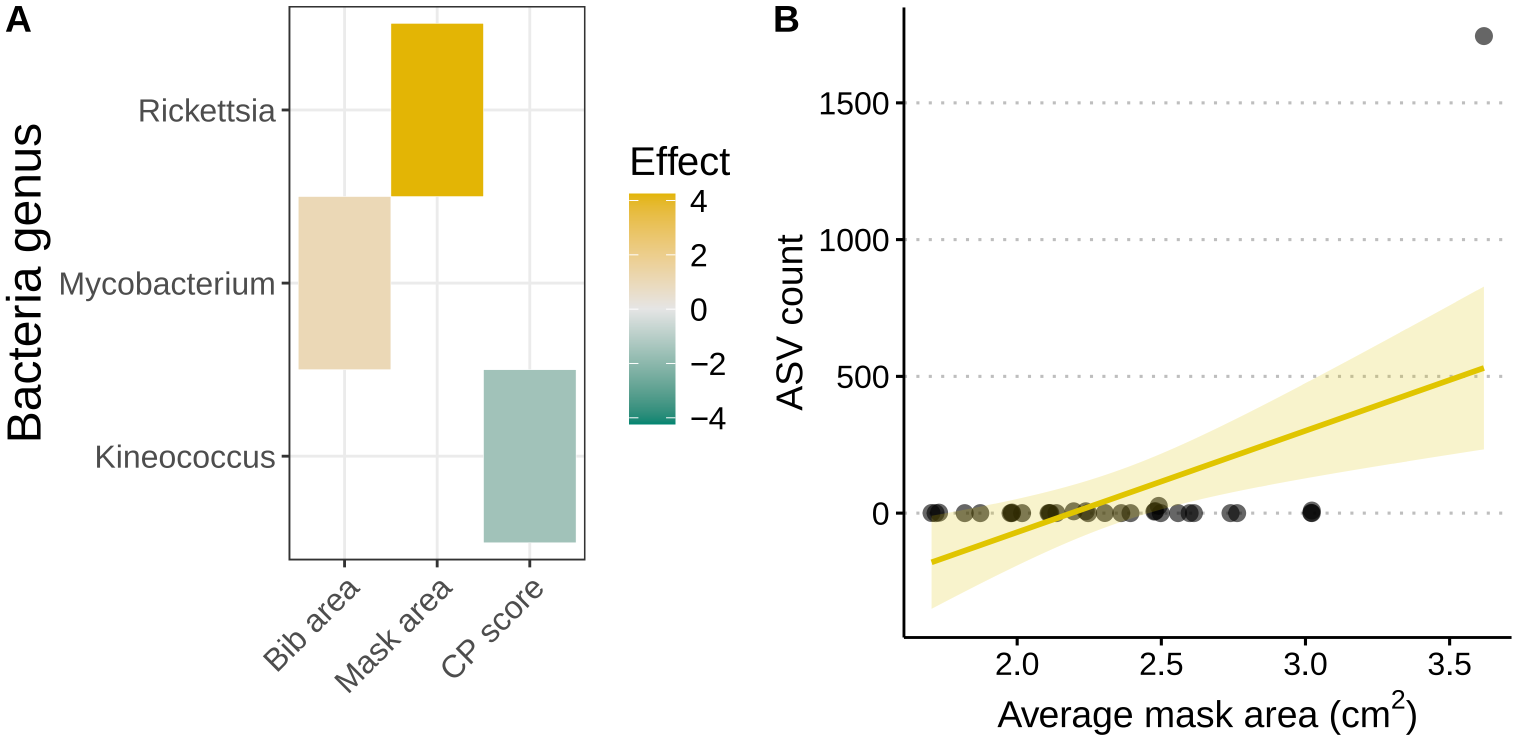
