## Supplemental Legends for "Colorful connections: pigment-based plumage and breeding condition are associated with gut microbiome variation in the Common Yellowthroat"

Figure S1. Example images of male Common Yellowthroats depicting variation in phenotypic traits, including (A) standardized photos against a 1cm checkered grid, (B) the spectrum of bib sizes (outlines only) visually and numerically, and (C) the spectrum of mask sizes visually and numerically.

Figure S2. Associations of significantly differentially abundant ASVs using the New York-only dataset. (A) Heatmap illustrating ASV genera that are most strongly (Q < 0.05) associated with host plumage features. Colors represent the strength and direction of the association (effect score = −log10(Q-value) × sign(coefficient)). Teal represents negative associations and gold represents positive associations. (B) Counts of *Rickettsia* (sum of congeneric ASV read counts) in relation to average mask area. Each point represents a sample and trend lines represent linear regression fits with 95% confidence intervals.

Table S1. Comparison of competing models (ΔAICc ≤ 2) for alpha diversity metrics using Akaike’s Information Criterion adjusted for small sample size (AICc). Each tab contains results for a specific diversity metric (e.g., Shannon, Chao1, and Faith’s phylogenetic diversity) and data subset (e.g., all samples or New York-only samples). The null model (i.e., intercept-only) is shown for comparison as well. Columns include: “model_id” to identify each candidate model with an alphanumeric code; “term” to indicate model predictors; “estimate” and “std_error” are model coefficients and associated standard errors for each term; “lower_85” and “upper_85” represent 85% confidence intervals; “K” is the number of parameters; “AICc” is the Akaike Information Criterion corrected for small sample size; “Delta_AICc” is the difference in AICc relative to the top-ranked model; and “AICcWt” is the model weight.

Table S2. Results from the top 10% of models as ranked by adjusted R^2^ for beta-diversity metrics. Each tab contains results for a specific diversity metric (e.g., Jaccard, Bray-Curtis, unweighted UniFrac, and weighted UniFrac) and data subset (e.g., all samples or New York-only samples). Columns include: “predictors” which lists the full set of predictors included in each model; “global_F” and “global_p” are the F-statistic and associated p-value for the overall permutation test; “globally_significant” indicates whether the model is statistically significant; “R2_adj” is the adjusted coefficient of determination; “predictor” indicates individual terms within the model; “marginal_F” and “marginal_p” are the F-statistic and p-value for each predictor (marginal tests); and “model_id” identifies each candidate model with a numeric code.

Table S3. Modelling results from differential abundance analyses at the ASV level, only displaying the associations for which the FDR-adjusted P-values (Q-values, “qval”) are below a threshold of 0.1. Each tab contains results for a specific data subset (e.g., all samples or New York-only samples). Each row represents a test between a particular ASV (“feature”) and a metadata variable (“metadata”). Other columns include the reference level (“value”), the estimated effect size (“coef”) and its standard error (“stderr”), the number of samples included in the model (“N”), the number of samples for which the feature was non-zero (“N.not.0”), the raw (“pval”) and adjusted p-values (“qval”).
